# Peptide Aggregation Induced Immunogenic Rupture (PAIIR)

**DOI:** 10.1101/2021.12.11.472230

**Authors:** Gokhan Gunay, Seren Hamsici, Gillian A. Lang, Mark L. Lang, Susan Kovats, Handan Acar

## Abstract

Under the influence of stress and membrane damage, cells undergo immunogenic cell death (ICD), which involves the release of damage associated molecular patterns (DAMPs), natural adjuvants for enhancing an immune response. In the presence of an antigen, released DAMPs can determine the type and magnitude of the immune response, and therefore the longevity and efficacy of an antigen-specific immunity. In the last decade, the immune response effect of ICD has been shown, yet there is no tool that can induce controlled ICD with predictable results, regardless of the cell type. We designed a peptide-based tool, called [II], for controlled damage to cell membrane to induce ICD and DAMPs release. Herein we describe a series of experiments that determine that the mechanism of action of [II] includes a caspase-dependent ICD and subsequent release of immune stimulating DAMPs, on various cell types. Moreover, we tested the hypothesis that controlled DAMP release via [II] in vivo was associated with enhancement of antigen-specific adaptive immunity with influenza hemagglutinin (HA) subunit vaccine. HA and [II] showed significantly higher HA specific IgG1 and IgG2a antibodies, compared to HA-only immunized mice, while the peptide itself did not elicit antibodies. In this paper, we demonstrate the first peptide-aggregation induced immunogenic rupture (PAIIR) approach as vaccine adjuvants for increasing both humoral and cellular immunity. In consideration of its ability to enhance IgG2a responses that are associated with heterosubtypic influenza virus protection, PAIIR is a promising adjuvant to promote universal protection upon influenza HA vaccination.

---

As a result of cell membrane damage or stress, cells release damage-associated molecular patterns (DAMPs) and die through immunogenic cell death (ICD), which has significant immune system enhancing effects. Several infectious pathogens and cancers deploy strategies to limit the emission of DAMPs and escape immunosurveillance. ^1^ Inducing and controlling localized ICD in the presence of an antigen has a significant potential for therapeutic and protective immunity. Released DAMPs with immunostimulatory effects show “adjuvant-like” properties and engage antigen presenting cells (APCs) for the stimulation of robust, long-lasting immunity. In cancer treatment, inducing ICD and DAMP release with chemotherapeutics and photothermal therapy substantially improves the antitumoral effect. ^2,3^ Similarly, engaging DAMP release enhances the antigen-specific immunity in vaccination strategies. ^4^ Despite this potential, the current ICD-inducing technologies are limited because they are effective only in specific applications, and demonstrate off-target toxicity. ^5,6^ Therefore, there is a significant clinical need for a tool that can induce controlled ICD and is effective across different cell types to increase the efficacy of current immunotherapy treatments.

Nanomaterials built by self-assembling peptide have wide application areas in medicine. Discovery of peptide sequences that assemble into nanostructures with desired properties and functions in a biological environment is a major challenge for the field. Editing the sequence of a protein that naturally assembles in the body can accelerate the discovery of various sequences, yet the harvested properties rule their functionalities. Establishing the application of a new peptide material based on its characterized properties is a common strategy. ^7^ Here we describe a different approach, in which we first determined the properties of the desired material, based on knowledge of proteins that induce immunogenic cell death (ICD), and then designed peptide sequences that can create similar effects. Regardless of the organism in which it occurs, the ICD process follows a similar trend: aggregation of cell membrane pore forming proteins (or peptides) cause damage to the cell membrane after activation, and this damage subsequently releases natural adjuvants called damage-associated molecular patterns (DAMPs), such as ATP, DNA, high-mobility group box 1 (HMGB-1), and heat shock protein 90 (HSP90).^8,9^ To elicit a similar aggregation kinetics with a small peptide sequence, we utilized the co-assembly of oppositely charged peptides (CoOP) strategy for this design. ^10^

CoOP strategy is based on a framework (i.e., the diphenylalanine (FF) domain and terminal charges) that defines the peptide-peptide orientation, and thus initiates the interactions of the peptides. However, the forces that affect the kinetics of the assembly and the properties of the final product are provided by the two amino acids of the substitution domain [XX] (we use the standard single-letter amino-acid codes throughout this work, with the CoOP pairs denoted in square brackets: [ ]). Our CoOP strategy provides quantitative correlations between changes in the amino acid sequence of the framework and properties of the assembly. ^10^ Hence, CoOP represents a uniquely powerful strategy to create small peptides with controllable aggregation profiles that can be used to design peptide-aggregation-induced immunogenic rupture (PAIIR) of the cell membrane and subsequent ICD.

Adjuvants induce antigen-specific antibodies to either tag the antigens on pathogens or infected cells for attack by effector immune cells (as in type-1 cellular response correlated to Th1) or neutralize antigen-carrying entities (as in type-2 humoral response correlated to the CD4+ T cells called T helper 2, Th2). ^4,11^ Most common adjuvants enhance type-2 response via Th2 and produce neutralizing antibodies, immunoglobulin G1 (IgG1). A typical Th1 response to a viral infection is characterized by recruitment of cytotoxic T lymphocytes (CTL), macrophages and NK cells, and associated with IgG2a or IgG2c formation in mice. APCs recognize DAMPs through their pattern recognition receptors (PRRs) and become activated, ^12^ create non-infectious local inflammation and enhance their ability to internalize and present antigens to activate the adaptive immune response. ^13^ For example, oil-in-water emulsion MF59 induced ATP release from muscle cells through intramuscular vaccination, and degradation of extracellular ATP reduced hemagglutinin (HA)-specific antibody formation when mice immunized with HA-influenza antigens. ^14^ However, ATP injection alone was not sufficient to induce a strong response, suggesting a synergistic effect that relies on other released DAMPs. ^14^ In a recent study, plasmid-encoded HMGB-1 increased the immunogenicity of a DNA vaccine, thus enhancing a specific immune response to the influenza nucleoprotein and hemagglutinin proteins, and improving the survival of mice against lethal mucosal heterosubtypic influenza challenge. ^15^ HSP90 enhances the cross-presentation of antigens by APCs.^16^ However, recombinant DAMPs have been ineffective adjuvants. ^17^ Therefore, technologies that induce release of DAMPs collectively for synergistic effects are an emerging frontier in immunotherapy.

Vaccination is critical for preventing the spread of respiratory viruses like influenza. However, current influenza vaccines have limited efficacy and cannot provide long-lasting and universal protection due to the high mutation rate in HA antigens.^18–20^ Induction of a strong cellular immunity against more conserved antigens of influenza is critical for universal protection, which relies on a Th1 related immune response. ^21^ However, vaccination with the influenza HA protein combined with the majority of the commercially available adjuvants induces an IgG1 response, limiting long-lasting universal protection. ^22^ Here, we hypothesize that, peptide-aggregation induced immunogenic rupture on cells can be utilized as a vaccine adjuvant. By mixing our new peptide tool with influenza HA proteins, we developed an influenza vaccine and studied the immune response (**Figure 1**). We demonstrate the use of the CoOP strategy to design peptide sequences with desired aggregation properties to compose PAIIR. We identify the mechanism of programmed ICD and demonstrate DAMP release induced by our designed peptides in multiple cell types. Upon co-administration, PAIIR enhances HA-specific IgG1 and IgG2 formation significantly, compared to immunization with HA alone. These results indicate that PAIIR induces Th1 and Th2 immune responses in mice. More importantly, we show that the designed peptide did not itself generate any specific antibody. These results highlight PAIIR as a promising new tool that can readily be injected, providing a simple, efficient, and safe method for enhancing immune response in vaccine applications.

**Figure 1:**
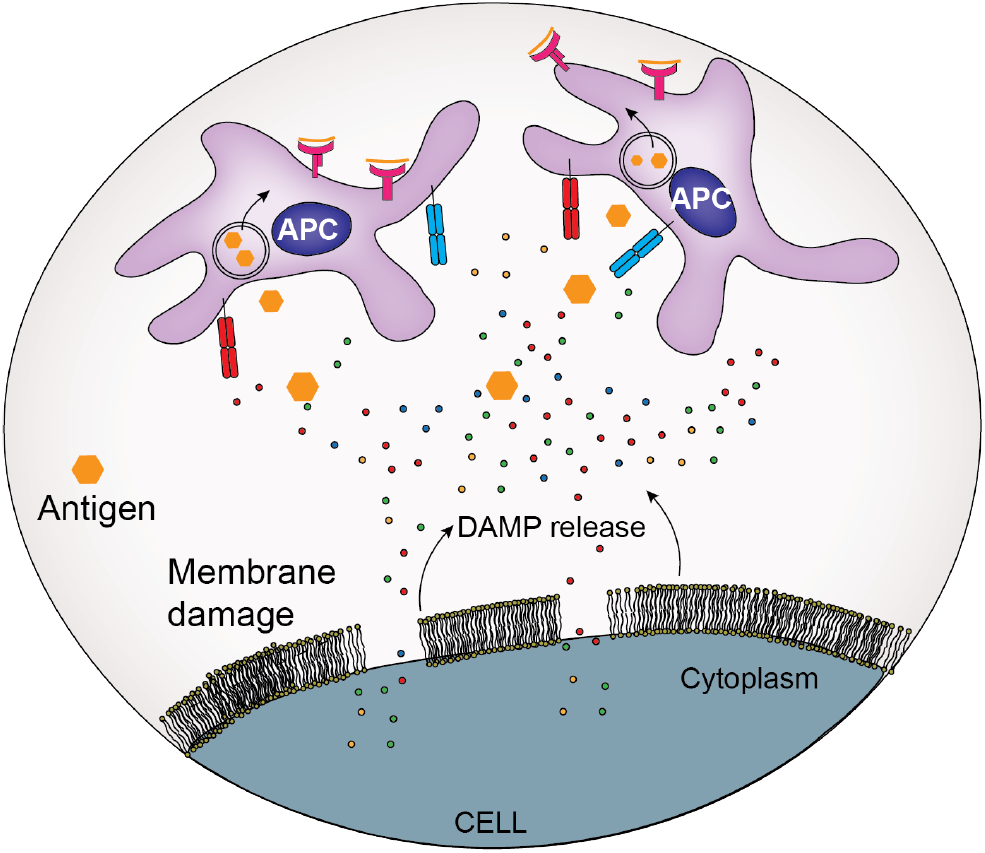
Cell membrane damage leads to release of DAMPs. These local DAMPs activate antigen presenting cells for the uptake, processing and presentation of antigens

Engineering the fine-tuned cellular aggregation of peptides requires well-characterized intermolecular interactions. The organization of peptide assemblies emerges from small changes in the amino acid sequence that provides non-covalent interactions, such as electrostatic forces, hydrogen bonds, hydrophobic effects, and aromatic stacking. ^23,24^ Understanding these interactions allows the rational design of peptides to encompass desired properties, such as aggregation on the cell and rupture of the membrane for DAMP release and ICD. For this purpose, we utilized the the CoOP strategy, where the the local charges promote electrostatic interactions between two oppositely charged amino acids; the anionic carboxylate of glutamate (E), and the cationic ammonium from Lysine (K)(**Figure 2A**). Electrostatic interactions are limited on the end of the peptides because of the Nterminal acetyl group and C-terminal amide group. ^10^The deliberately short, charged design enables the internalization of CoOP-based peptides into cells. ^25^ Design studies on multidomain peptides revealed the importance of hydrophobic localization in the core of the structures. ^26^ In particular, the interplay between the backbone hydrogen bonding and hydrophobic interactions among amino acid side-chains is crucial to the formation of fibrillar structures. ^27^ Further simplistic modifications of FF peptides (e.g., introducing aromatic capping groups ^28^ and co-assembly with hydrophobic amino acids ^29^) provided structural diversity in self-assembly products and control over the aggregation kinetics. Therefore, to change the aggregation kinetics, we studied the hydrophobic amino acids in the substitution domain (“[XX]”), with increasing hydrophobicity indices; [AA] *<* [VV] *<* [WW] *<* [LL] *<* [II] (**Figure 2A**).

**Figure 2:**
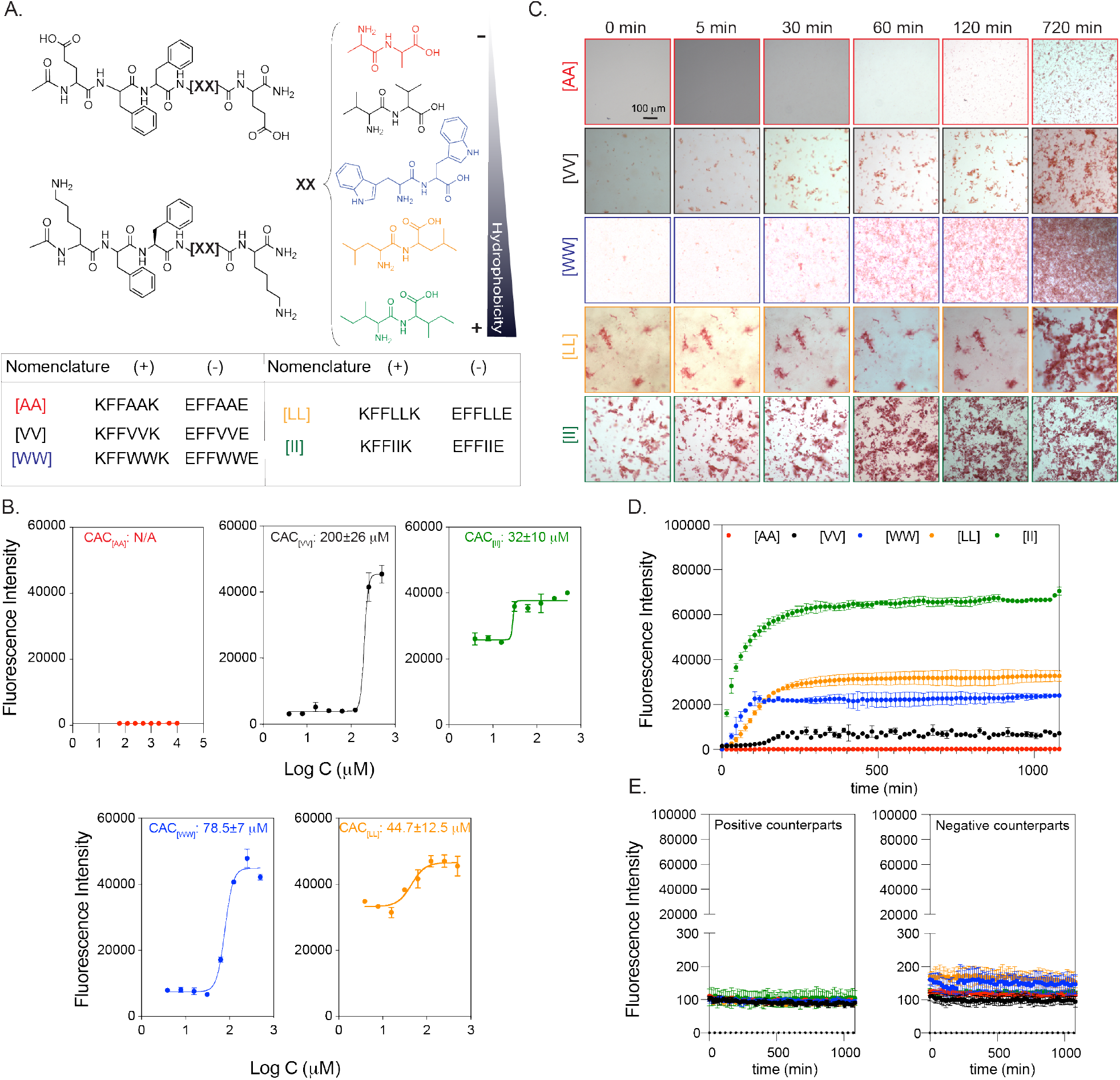
Peptide groups and aggregation kinetics. (A) CoOP groups and individual counterparts used in this study. (B) CAC of co-assembled peptides with DPH. (C) Macroscopic CoOP aggregates incubated with Congo Red. (D) Determination of aggregation kinetics of CoOPs. (E) and individual counterparts by ThT assay.Data are representative of three experiments.

The critical aggregation concentrations (CAC), i.e., the minimum concentration needed for coassembly of the peptides, are measured with DPH ((1,6-diphenyl-1,3,5-hexatriene)), which becomes fluorescent when located in aggregates with hydrophobic cores. ^30^ The CAC values of [VV]=200±26 *μ*m, [WW]=78.5±7 *μ*m, [LL]=44.7±12.5 *μ*m, and [II]=32±10 *μ*m followed the same trend as the hydrophobicity of the amino acids at their core (**Figure 2B**). The identified CAC values were correlated to our previous study. ^10^. No fluorescence was observed for [AA] even at 10 mm, indicating no aggregation; this situation therefore represents an excellent control for studying the effects of peptide aggregation, as the same concentration of peptides with similar sequence and charges will remain as free components in solution. The aggregation kinetics of the peptide pairs were measured with Congo Red and Thioflavin T (ThT) assays. Congo Red staining has been used to identify amyloid fibrils in vitro and in tissue sections, emits red light due to the binding of *β*-sheet rich domains, leading to a red shift in its absorbance peak from 490 to 512 nm. ^31^ We monitored the Congo red for 720 min (12 h) (**Figure 2C**) at 0.5 mm peptide concentration. [AA] precipitation appeared after 720 min; these are not likely to be ordered aggregates with hydrophobic cores because no DPH fluorescence was observed at this concentration. [II] aggregated instantaneously (0 min), while [LL] aggregates were deposited initially and became stable until 720 min. [WW] and [VV] did not show any initial aggregates but deposited after 30 min and 120 min, respectively. Despite ease of visualization with red light, Congo Red is not as sensitive as ThT in terms of understanding aggregation kinetics since fluorescence methods (rather than absorption) are preferred for high-sensitivity detection. ^32^ Therefore, we use Congo Red as a visualization method, with ThT used as for quantitative temporal measurements of oppositely charged pairs and each counterpart individually for 1200 min (20 h). To determine the time needed for equilibrium assembly of CoOPs, we applied curve fitting to the ThT analysis (**Figure 2D**). [AA] did not assemble, and thus did not show any fluorescence in ThT. For the remaining peptide pairs, the time to reach equilibrium assembly was calculated as [VV] (660 min) ¿ [WW] (675 min) ¿ [LL] (630 min) ¿ [II] (510 min). Individual counterparts did not produce any fluorescence signal (**Figure 2E**). We observed that the like-charged groups alone did not aggregate, highlighting the importance of electrostatic attractions in the aggregation process (**Figure 2D**). Among the studied peptides, [II] showed not only the fastest aggregation but also the highest fluorescence intensity, indicating that these peptides stacked in a more well-ordered structure than the other CoOPs. These results indicate that control over peptide aggregation is achieved by changing the hydrophobicity of amino acids in the substitution domain.

**Table 1:** The relationship of side chain properties with CAC and equilibrium time for co-assembly of CoOPs

|  | [AA] | [VV] | [WW] | [LL] | [II] |
| --- | --- | --- | --- | --- | --- |
| CAC [$\mu\text{M}$] | N/A | 200±26 | 78.5±7 | 44.7±12.5 | 32±10 |
| Equilibrium reaching time [min] | N/A | 690 | 675 | 630 | 510 |
| Sidechain hydrophobicity<sup>33</sup> | 41 | 76 | 97 | 97 | 99 |
| Hydrophobic/hydrophilic surface area<sup>34</sup> | 1.5 | 3.1 | 2.3 | 3.6 | 3.9 |
| t1/2 (half time, min) | N/A | 96.3±10.8 | 33.6±1.9 | 79.4±3.6 | 53.45±1.82 |

We used the human ovarian cancer cell line OVCAR-8 as our initial model to identify the effect of the peptides on cells. At 0.5 mm (i.e., above the overall CAC), the individual peptides mixed on the cells by adding first negative, then positive counterparts promptly. The viability of the cells was measured for 6h through live-dead imaging (**Figure 3A**). Among the peptide groups studied, [AA], [VV] and [LL] did not affect cell viability in the measured time frame (**Figure 3A**). Only [WW] and [II] caused cell death compared to control group. To understand the effect of peptide aggregates versus peptides alone on the cell death identified with [WW] and [II], we performed a live/dead assay with individual peptides at 0.5 mm after 6h (**Figure 3A**). Despite the dramatic cell death observed with the [II] mixture at this concentration, the individual [II] peptides did not induce any cell death. Similarly, the negative [WW] (EFFWWE) peptide alone did not show any cell death, yet the positive [WW] (KFFWWK) peptide induced cell death (as indicated by red stained cells). Positive charged peptides containing W are known to be promiscuous residues for membrane damaging peptides, possibly because of the “anchoring” role of W; W is abundant in membrane proteins, particularly near the lipid–water interface. ^35,36^ Furthermore, as analyzed by zeta-potential, individual peptides showed the expected overall charges at pH 7.0 ((pKa of (-COO^-^) of E is 4.25 and (-NH_3_^+^) of K is 10.53) (**Figure 3B, inset)**. In an aqueous solution, the [II] aggregates acquire a neutral charge in 5 min, while [WW] showed a slight positive charge after 30 min, possibly due to incomplete assembly (**Figure 3B)**.

**Figure 3:**
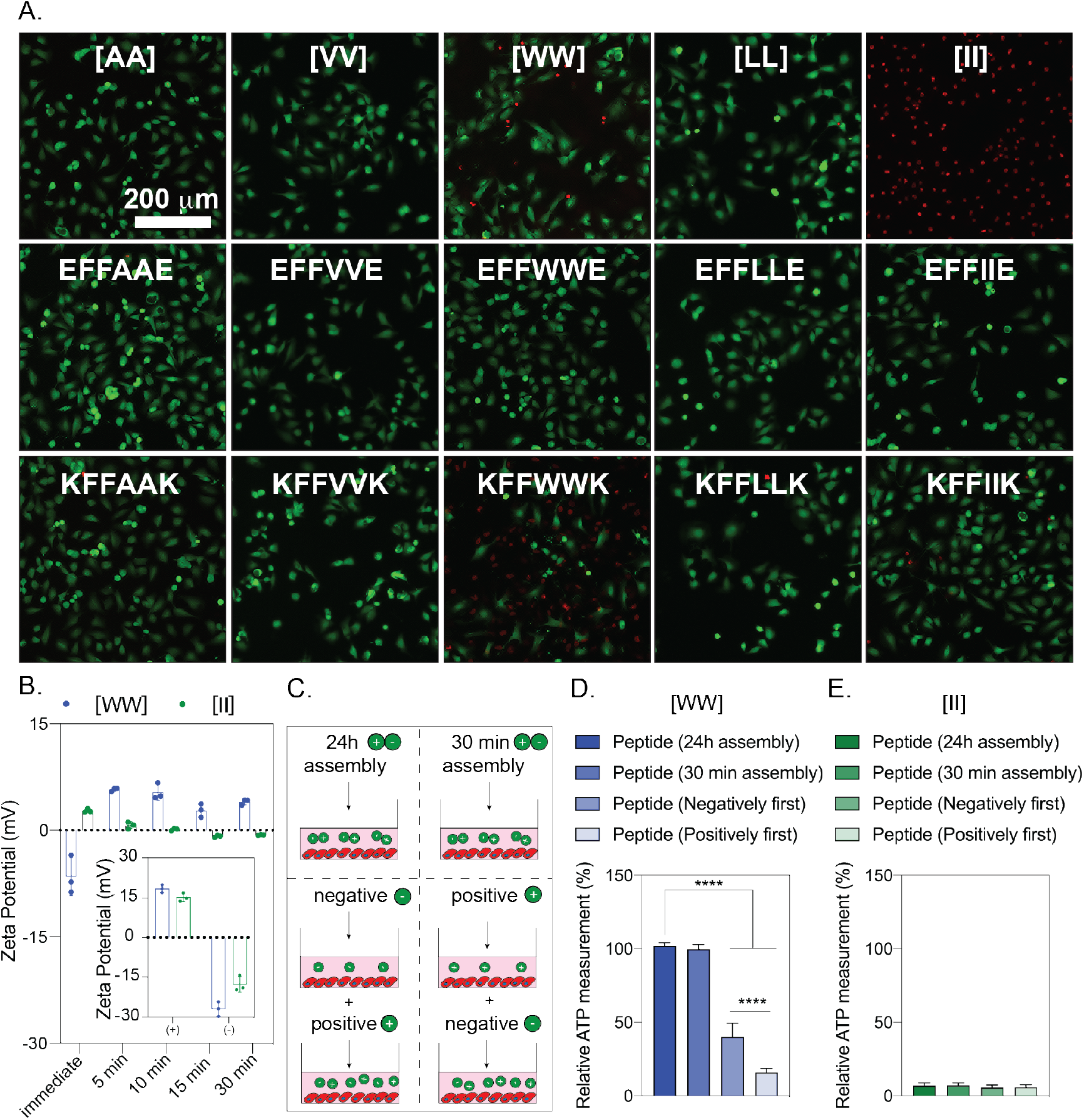
Cytotoxicity of peptides to OVCAR-8 cells. Negative peptides were added to the cells followed by the addition of corresponding positive peptides. (A) Live-dead images of OVCAR8 cells treated with peptide combinations and corresponding individual counterparts at 6h (peptide concentration is 0.5 mm). (B) Zeta potential of individual and combined peptides. (C) Illustration of peptide preparation for [WW] and [II]. The effect of peptide preparation on cell viability for [WW] (D) and [II] (E). Scale bar is 200 *μm*. Data are representative of at least three experiments. Statistical analysis via one-way ANOVA test, data are mean ±*SD;* [* * **]p¡0.0001.

Given the effects of [II] and [WW] on cell viability, it is interesting to examine both the order in which the positive and negative peptide counterparts are added and the effects of pre-mixing the peptides (i.e., the introduction of pre-aggregated peptides). We therefore changed the order of addition (i.e. positive first or negative first) and pre-mixed peptide components for 30 min or 24h before addition onto the cells (**Figure 3C**). Pre-mixing the peptides before addition to the cells completely abolished the cytotoxicity induced by [WW] (**Figure 3D**), indicating that the initial cytotoxicity was due to the positive charge of KFFWWK that is neutralized upon mixing and aggregation with the counterpart (**Figure 2B**). Addition of KFFWWK first induced significantly higher cytotoxicity ([****]*p <* 0.0001) compared to the initial addition of negative peptide, indicating that the cytotoxicity is due to the positive charge of KFFWWK (**Figure 3D**). Nevertheless, none of the studied conditions affected the cytotoxicity levels for [II] (**Figure 3E**), highlighting that [II] is the only peptide pair among those studied that induces cell death through aggregation alone.

Clearly, the individual peptides of the designed [II] are not triggering any cell death pathway (**Figure 3A**), which indicates that the aggregation of the [II] peptide causes the cell membrane damage. There are other peptides derived from natural proteins that induce ICD through perturbation of both cell and mitochondrial membranes. ^37–39^ However, these peptides targets Bcl-2 family proteins to create mitochondrial damage. ^40,41^ The oncogenic mutations in the Bcl-2 proteins and known drug resistance against therapeutics targeting them, diminishes the effectiveness of these peptides. ^42,43^

As aggregation-induced cytotoxicity was only observed with [II], we analyzed the mechanism by which it induces cell death. We mixed EFFIIE and KFFIIK for 30 min prior to administration on cells. To understand cell membrane damage (necrosis), we used propidium iodide (PI), a dye that does not permeate intact cell membranes. ^44^ **Figure 4A** shows cells treated with three different concentrations of [II]: 0.25 mm, 0.5 mm and 1mm for 1-6 and 24h. Increasing the concentration to 1mm resulted in faster membrane damage, while decreasing the concentration to 0.25 mm (although still higher than the [II] CAC of 38 *μ*m) did not have sufficient aggregation to trigger cytotoxicity in this time frame. The time and concentration dependence of [II] cytotoxicity is likely to be a result of faster aggregation in higher concentrations. The relative viability of the cells was measured by intracellular ATP content, indicated by the ability of [II] at 0.5 mm to induce *>*85% of cell death in 6h (**Figure 4B**). Cell swelling and nuclear condensation, hallmarks of necrotic cell death, were observed in the morphology of [II] incubated cells (**Figure 4C and D**). ^45^ These results show that [II] induced cell death can be controlled through time and concentration. To visualize the aggregation, positive charged [II] (KFFIIK) peptide was labeled with FITC. We monitored the internalization and aggregation of KFFIIK through fluorescence microscopy. KFFIIK alone was internalized into the cells starting at 1h (**Figure 4E**), and cellular morphology did not changed over time as KFFIIK alone is not cytotoxic. When KFFIIK was introduced with its negatively charged counterpart, EFFIIE, the peptide-aggregation (more localized green KFFIIK) was apparent around and within the cells starting at 3h (**Figure 4F**). The peptide-aggregation induced nuclear area shrinking and loss of integrity of the cells (observed through absence of actin fibers), indicating [II] aggregation induced cell membrane damage and death.

**Figure 4:**
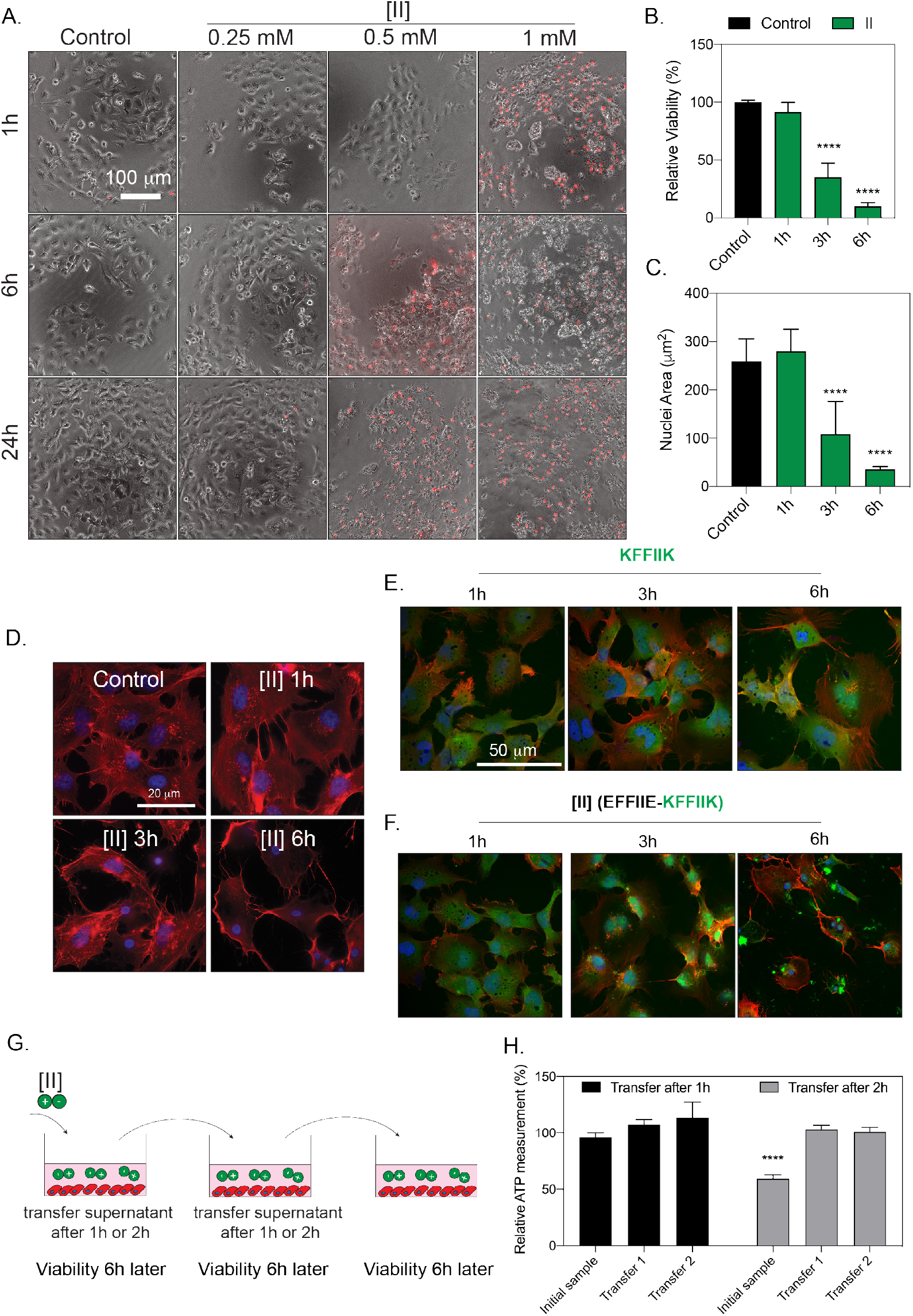
[II] induces membrane rupture. (A) Time and concentration dependent cytotoxicity of [II] analyzed through PI uptake (scale bar is 100 *μm*). Time dependent viability measurement (B), and time dependent nuclear size analysis (C). Actin cytoskeleton staining of [II] treated OVCAR-8 cells at 1,3 and 6h, scale bar is 20 *μm* (D). Internalization of FITC-KFFIIK (E) and FITC-[II] for 1,3 and 6h (F), scale bar is 100 *μm*. Experimental plan of [II] transfer and viability experiment (G). Effect of [II] transfer on cell viability at 6h (H). Data are representative of at least three experiments. Statistical analysis was done with one-way ANOVA test, data are mean ±*SD*, [* * **]p¡0.0001

Observation of cell death beyond the desired area (i.e., off-site cytotoxicity) is a major obstacle for existing therapeutic applications. To estimate the off-site cytotoxicity of our approach, we incubated cells with [II] for 1h or 2h followed by the transfer of supernatant into another well with fresh cells, and measured viability after 6h (**Figure 4G**). We observed cell death only in the initial sample after 2h (**Figure 4H**), suggesting that at this concentration, [II] does not induce cytotoxicity in 1h (in line with the viability results **Figure 4B**). Also, subsequent transfer of the supernatant of cells incubated with [II] for 1h does not induce cytotoxicity, possibly due to aggregation having started within the cells and thus insufficient [II] transfer to the next well.

DAMP release through cell membrane rupture is a hallmark of ICD. To quantify cell membrane damage we measured the release of lactate dehydrogenase (LDH), a cytoplasmic enzyme released into the extracellular matrix upon membrane damage. ^46^ LDH release became significant 3h after incubation with [II] (**Figure 5A**), in agreement with the reduction of cell viability (**Figure 4B**). As expected, significant DAMP release followed the cell membrane damage, such as the presence of extracellular ATP ([****]*p <* 0.0001) after 3h (**Figure 5B**), which was correlated with LDH release. However, the amount of extracellular ATP is depleted in 6h due to cytotoxicity *>* 85% (i.e., few live cells remain to release ATP upon ICD) as well as rapid extracellular hydrolysis of ATP. ^47^ Similarly, the release of other DAMPs (specifically HMGB-1 and HSP90) was also detected in the extracellular environment of [II] treated cells at 6h via western blotting (**Figure 5C**). Our DAMP release profile results highlight the immunogenicity of the cell death induced by [II] peptide pair aggregation. *In other words, peptide-aggregation induced immune rupture (PAIIR) is demonstrated with the [II] pair*.

**Figure 5:**
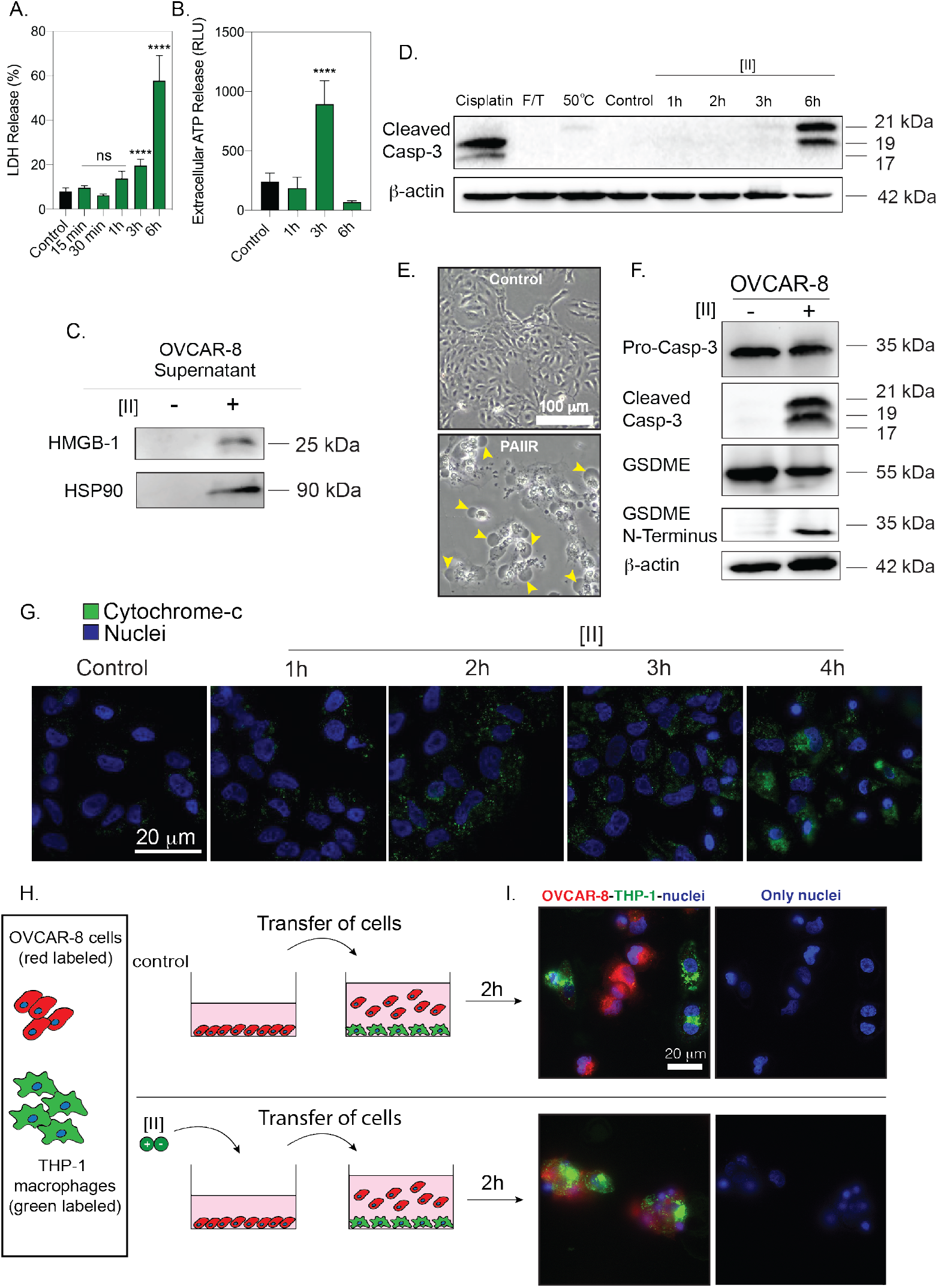
[II] induces DAMP release and secondary pyroptosis. Time dependent LDH release (A). Time dependent extracellular ATP release (B). Western blot analysis of HMGB-1 and HSP90 release at 6h (C). Caspase-3 cleavage after treatment with Cisplatin, F/T, heat treatment and 1-2-3 and 6h of [II] treatment (D). Phase contrast images of control and [II] treated OVCAR8 cells at 6h, yellow arrows indicate bubbles emerging from pyroptotic cells (E). Caspase-3 and GSDME cleavage of OVCAR-8 cells with and without treatment with [II] (F). Time dependent cytochrome-c release into cytoplasm from OVCAR-8 cells (G). Scale bar is 100 *μm*. Schematic illustration of the phagocytosis assay (H). Phagocytosis of OVCAR-8 cells by THP-1 differentiated macrophages after 6h of [II] and control treatment (I) Data are representative of at least three experiments..

Regulated cell death via an inducer is concentration and time-dependent, and therefore controllable. Unregulated cell death, on the other hand, cannot be controlled to the same extent as it mainly results from mechanical forces or temperature. DAMP release from regulated cell death is prolonged, which is essential for the desired immune response, while unregulated cell death essentially releases an instantaneous burst of DAMPs. Therefore, regulated ICD is a desired for a controlled immune response. Regulated necrotic cell deaths are caspase-dependent^48,49^, as is caspase-3 cleavage in regulated apoptosis. ^50^ To identify whether [II]-induced cell death is regulated, we analyzed the cleavage of caspase-3, a known mediator of apoptosis and GSDME mediated pyroptosis (a regulated necrosis). ^2^ We compared the caspase-3 cleavage among different cell death modalities; as an apoptosis control, we used cisplatin - a known apoptosis inducer in OVCAR-8 cell line. ^51^ We also used heat treatment to induce multiple cell death modalities, including necrosis, apoptosis, and necroptosis. ^52^ Additionally, we applied a freeze/thaw (F/T) method, a common technique to induce unregulated necrosis. ^53^ (**Figure 5D**) shows western blotting of caspase-3 cleavage upon these treatments. Cisplatin showed caspase-3 cleavage, which is expected in apoptotic cell death. Conversely, F/T did not show any caspase-3 cleavage. Observation of cleaved caspase-3 in [II] induced cell death indicates regulated necrosis (e.g., GSDME mediated pyroptosis).^54^ Indeed, pyroptosis can be identified via cleavage of caspase-3 and GSDME and a ballooning morphology of the cell membrane. ^55^ As cell membrane damage is already identified, we examined the morphological features of [II] treated cells and observed ballooning of the plasma membrane (**Figure 5E**), yellow arrows). At the molecular level, cleavage of caspase-3 activates GSDME cleavage, then the N-termini of the cleaved GSDME oligomerize and induce membrane pore formation. ^8^ Western blot analysis showed that OVCAR-8 cells express GSDME and treatment with [II] results in its cleavage at 6h (**Figure 5F**). These results show that [II] treated cells have the morphological and molecular features of GSDME-mediated pyroptosis, a regulated ICD.

Given these results, we then studied the events upstream of caspase-3 activation. The majority of caspase activation within mammalian cells is initiated through cytochrome c, a protein normally found in mitochondria. ^56^. Released cytochrome c initiates the activation of caspase-3^57^ and the downstream pathway of cytochrome c release is not dependent on the functions of BAX or BAK proteins. ^41^ Furthermore, cytoplasmic cytochrome c release has been shown to lead to the induction of GSDME-mediated pyroptosis in cells that express GSDME.^55^ To understand the origin of caspase-3 cleavage upon [II] treatment, we stained cytoplasmic cytochrome c to analyze mitochondrial damage as shown before. ^58^ Time-dependent imaging showed that cytochrome c levels increased over time in the cytoplasm of OVCAR-8 cells (**Figure 5G**), specifically at 4h, explaining the abundance of cleaved caspase-3 at 6h (**Figure 5D**). These results show that [II] peptide induces mitochondrial membrane damage, leading to cytochrome c release into the cytoplasm; this release activates caspase-3 cleavage, and initiates GSDME-mediated pyroptosis.

As [II] induced cells showed regulated ICD and DAMP release, which enhanced the phagocytosis of APCs, ^53^ we checked whether [II] treated cells are phagocytosed by macrophages. THP-1 differentiated macrophages are commonly utilized for in vitro phagocytosis experiments^59^. We treated red-labeled OVCAR-8 cells 6h with [II] and co-cultured them with green-labeled THP-1 macrophages for 2h. We observed phagocytosis of [II] treated cells by the macrophages, yet control cells were not phagocytosed(**Figure 5I**).

In cells where GSDME is not expressed, apoptosis is observed instead of pyroptosis, which is immunologically silent.^60^ To identify whether [II] induced ICD require GSDME expression, we tested various cell lines that have different GSDME expression profiles; B16F10 and Panc02 cells lines showed no GSDDME expression, while 3T3 fibroblasts had low expression levels compared to OVCAR-8 (**Figure 6A**). The incubation of [II] (0.5 mm) with all these cell lines under the same conditions showed a similar cell membrane rupture effect, as indicated by PI uptake (**Figure 6B**). [II] initiated the extracellular release of DAMPs; HMGB-1 and HSP90 in 6h (**Figure 6C**). Bubbling cell membrane structures were not observed in the cells that lacked GSDME (**Figure 6A**) suggesting that pyroptosis did not occur. These results show that [II] induced ICD is not dependent on GSDME, although it also has a GSDME-regulated secondary mechanism. Expression of GSDME was linked to anti-tumor immunity and tumor suppression^60^. However, GSDME is expressed at low levels in most cancer types ^61^ due to epigenetic silencing through methylation^62^.

**Figure 6:**
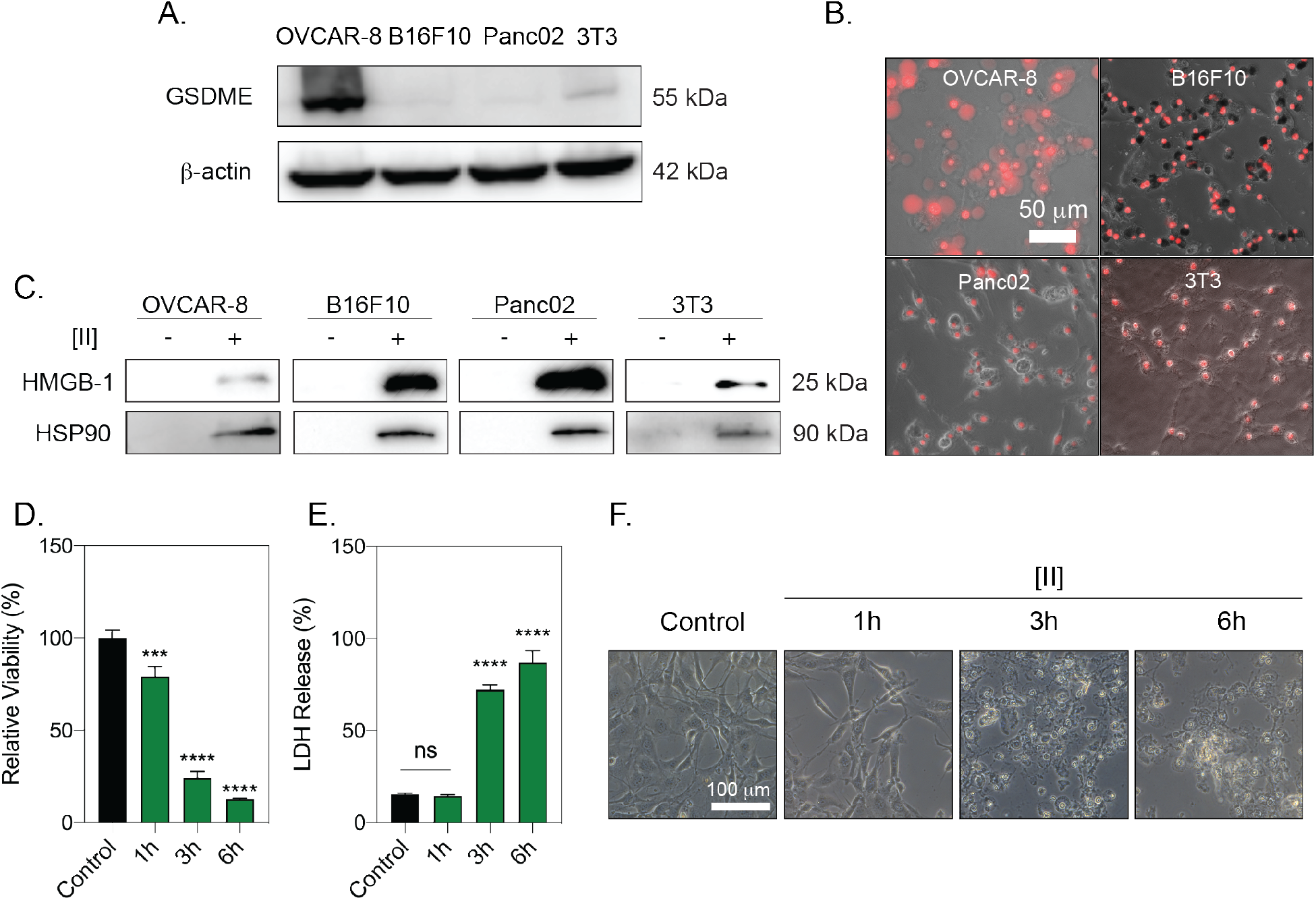
[II] induced membrane rupture and DAMP release is not cell type specific. GSDME analysis of OVCAR-8, B16F10, Panc02 and 3T3 cells (A). PI uptake at 6h of [II] treatment on cells (B). HMGB-1 and HSP90 release to the supernatant (SN) at 6h (C). Time dependent viability measurement (D). Time dependent LDH release (E). Phase contrast images of [II] treated fibroblasts at 1, 3 and 6h (F). Data are representative of at least three experiments. Statistical analysis was done by one-way ANOVA test, data shown are mean SD,[****]*p <* 0.0001, [***]*p <* 0.001

One of the cell types we analyzed was fibroblasts, an abundant cell type in the subcutaneous layer. ^63^ Stimulation of DAMP release from fibroblasts would act as an adjuvant following subcutaneous vaccinations. Therefore, we quantified the [II] induced ICD of 3T3 fibroblasts with time dependent cell death and LDH release ((**Figure 6D, E and F**), and the effect of [II] on fibroblasts was similar to that on OVCAR8.

We tested the hypothesis that the DAMP release caused by [II] and documented in **Figure 1 to 3**) was associated with enhanced IgG responses to influenza hemagglutinin (HA) (**Figure 7A**). Balb/c mice were immunized with PBS vehicle, recombinant HA, [II], or HA plus [II]. After 60 days, mice received booster vaccines identical to the initial vaccine dose. Mice were bled at 2 time points after the primary immunization (days 14 and 28) and once after the booster vaccine (day 75). HA-specific IgG1 could not be detected following administration of PBS vehicle or [II] alone (**Figure 7B**). HA-specific IgG1 was detected and increased with time since the primary immunization and was increased over 10 fold by the booster vaccine. At each of these time points, inclusion of [II] in the formulation caused a 2-4 fold and significant increase in HA-specific IgG1 titers. A similar pattern was observed for the less abundant IgG2a subclass, where [II] HA exerted a strong and significant adjuvant effect on the HA-specific IgG2a titer (**Figure 7C**). By day 28, [II] increased HA-specific IgG2a titers 7.3 fold and 12.4 fold by day 75. Consistent with the observation that [II] alone did not induce HA-specific IgG1 or IgG2a responses, ELISA plates were also coated with [II] (**Figure 7D**). [II]-specific IgG1 and IgG2a could not be detected. These results therefore demonstrate that [II] exerts a modest adjuvant effect on HA-specific IgG1 responses and a strong adjuvant effect on HA-specific IgG2a responses without inducing an IgG response to itself. These results are consistent with [II]-induced ICD and DAMP-release enhancing Th1 and Th2 Ab responses. Our observations are of particular significance because murine IgG2a was found to show higher cross-protection (heterosubtypic) against lethal challenge with different influenza virus strain. ^11,64,65^ Given the need for a universal influenza vaccine, any adjuvant that can boost the Th1 IgG response could contribute to achieving that goal.

**Figure 7:**
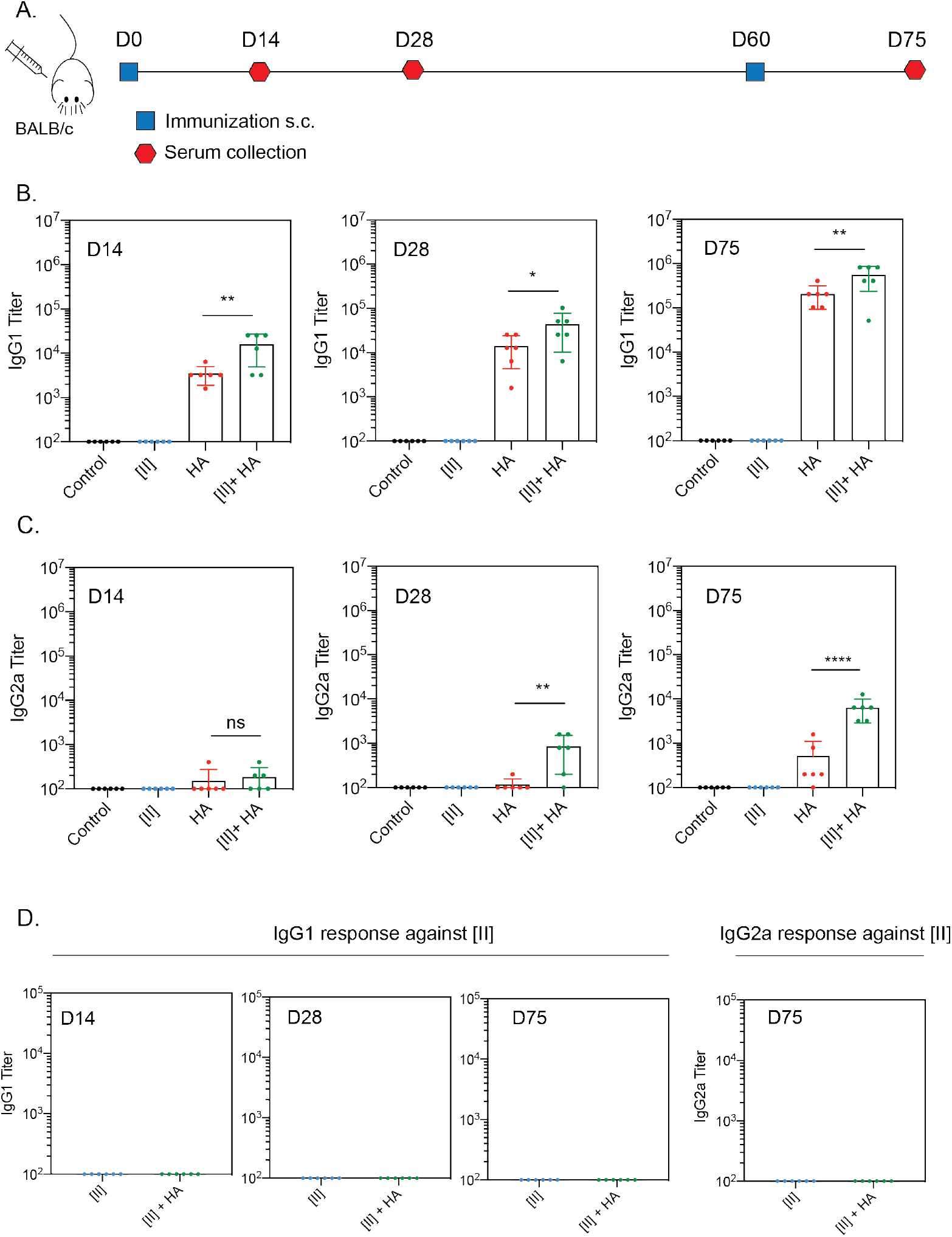
PAIIR induces influenza-antigen specific antibody formation. Timeline and the plan of the vaccination study: BALB/c mice were vaccinated at D0 followed by a booster at D60, serum was collected at D14, 28 and 75 (A). Antigen-specific serum IgG1 titers at D14, 28 and 75 (B). Serum IgG2a titers at D14, 28 and 75 (C).Serum IgG1 and IgG2a titers against [II] peptide at D14, D28 and D75 (D). Statistical analysis was done by one-way analysis of variance (ANOVA) with Tukey’s multiple-comparison test, data shown are mean SD [****]*p <* 0.0001, [**]*p <* 0.01, [*]*p <* 0.1 n=6 mice.

The lack of a [II]-specific IgG response is informative. The Major Histocompatibility Complex (MHC) class II is expressed by professional APCs. Loading of MHC molecules with self and foreign antigen-derived peptides governs immunological tolerance and the immune response to pathogens, respectively. The peptide-binding groove of MHC-II accommodates peptides of 13-25 amino acids. ^66^ The B cell antigen receptor (BCR) recognizes linear and conformational epitopes consisting of peptide sequences, or other macromolecules. Because [II] is a peptide-based adjuvant, the possibility of MHC-II presentation or recognition by the BCR was considered. Our results indicate that it is unlikely that the hexameric [II] peptide can bind MHC-II or be recognized by the BCR.

Interest in peptide-based materials has flourished in health, energy, materials science, and national security. ^23^ By advancing discovery of peptide domains with unique intermolecular interactions, design of peptides with desired properties in appropriate conditions can be achievable. Inspired by the mechanism of natural membrane rupturing proteins that aggregate and induce ICD, in this study we designed a new peptide-based tool, [II], via the CoOP strategy.

Understanding and controlling ICD is an essential tool that plays a paramount role in advancing cancer immunotherapy and vaccine development. We demonstrated the controlled ICD mechanism of [II] in vitro with various cell lines and observed IgG profiles secondary to Th1 and Th2 (i.e., cellular and humoral) immune responses in mice vaccinated with the [II] and influenza HA subunit. Peptide-aggregate induced immune rupture (PAIIR) induced by [II] pairs (“pa[II]r”) has tremendous potential to advance healthcare and basic science applications based on immune system modulation.

## 1 Experimental Section

### Materials

9-fluorenylmethoxycarbonyl (Fmoc) protected amino acids, [4-[*α*-(2’,4’-dimethoxyphenyl) Fmoc aminomethyl] phenoxy] acetamidonorleucyl-MBHA resin (Rink amide MBHA resin), Oxyma, N,N’- Diisopropylcarbodiimide (DIC), Trifluoroacetic acid (TFA), piperidine, dimethylformamide (DMF), dichloromethane (DCM) were purchased from Gyros Protein Technologies. Triisopropylsilane, Acetic anhydride, Congo Red dye and pyrene were purchased from Sigma-Aldrich. Deionized water (resistance of 18.2 MΩ.cm) was used during the experiments.

### Cell Culture and Reagents

Epithelial ovarian cancer cell line OVCAR-8 (NCI-Vial Designation 0507715), B16-F10, Panc02, 3T3 and THP-1 cell lines were cultured in humidified incubator at 37 ºC supplied with 5% CO2. B16-F10 and Panc02 cell lines were gifts from Dr. Wei R. Chen, University of Oklahoma. THP-1 cell line was gift from Dr. Stefan Wilhelm, University of Oklahoma. OVCAR-8, THP-1, Panc02 cells a were cultured in Roswell Park Memorial Institute (RPMI) media (SIGMA R8758) and B16F10 and 3T3 cells were cultured in Dulbecco’s Modified Eagle Medium (DMEM) (Sigma, D6429), supplemented with 10% FBS (Hyclone SH30910.03) and 1% antibiotics; penicillin (100 U/mL), and streptomycin (100 *μ*g/mL) (Thermo Fisher 15240062) according to the manufacturer’s instructions. To culture cells T75 flask (TPP 90076) were used and cells were passaged upon 85% confluency by using trypsin (Sigma 59418C). Media was changed every 2 days.

### Synthesis and Characterization of FF peptides

FF peptides (KFFAAK, EFFAAE, KFFWWK, EFFWWE, KFFIIK, and EFFIIE) were purchased from Biomatik (Biomatik Corporation, Canada) with higher than 95% purity. EFFVVE, EFFLLE, KFFVVK and EFFVVE peptides were synthesized via solid phase peptide synthesis with PreludeX automatic peptide synthesizer (Protein Technologies, Inc., Tucson, AZ). Peptides were prepared on a 0.2 scale by repeated amino acid couplings using Fmoc protected amino acid (3 eq.),DIC (7.5 eq.) and Oxyma (7.5 eq.). BHA Rink Amide resin was used as solid support to construct the peptides. Fmoc protected amino acids except the final residue were removed through treatment with 20% piperidine/DMF solution for 10 min (twice) at 50°C. Cleavage of the peptides from resin and deprotection of acid labile protected amino acids were carried out with a mixture of TFA/TIS/water in a ratio of 95:2.5:2.5 for 2.5 h. Excess TFA and organic solvents were removed by solvent evaporator and remaining peptide was precipitated using diethyl ether at 20°C overnight. The centrifuged white peptide precipitate was dissolved in water and frozen at -80°C overnight, and then lyophilized (Labconco Freezone, 12 L) for one or two days. Peptides were purified with preparative HPLC (Agilent 1260) and identified by Shimadzu LCMS-2020. Agilent ZORBAX 300 SB-C18 (9.4 × 250 mm) and Alltech Pro-sphere HP C4 300A 5u (250 mm x 4.6 mm) with a mo-bile phase of water/acetonitrile mixture (0.1% am-monium hydroxide) used for negatively charged peptides; water/acetonitrile mixture (0.1% formic acid) was used for positively charged peptides. All peptides tested with a purity *>*95%. HPLC run started with 100% water for 3 min, followed by a gradient increase in acetonitrile from 0% to 60% over 30 min, followed 100% acetonitrile for 3 min and finally 100% water for 3 min. Flow rate is 1.5 mL/min and injection volume are 10 *μ*L.

### Peptide aggregation analysis

DPH assay was performed to understand the CAC. Peptides (except [AA]) were prepared in PBS starting from concentration of 500 *μ*m to 3.9 *μ*m with serial dilution. [AA] was prepared from 10 mm to 15.6 *μ*m to check the aggregation behavior even in higher concentrations. First, negatively charged peptide (48 *μL*) was put to the black 96 wellplate, then 4 *μL* (from 4 *μ*m in PBS) DPH was added and finally positively charged peptide (48 *μL*) was added to each well. The solutions were incubated at 37°C for an hour. After the incubation, fluorescence intensity was collected immediately with Ex:360±40 nm, Em: 460±40 nm with with BioTek Neo2SM microplate reader. For visualizing aggregations deposited on the well plate, we performed Congo Red assay. Peptides having 0.5 mm concentration and equal volume (48 *μL* each) were prepared in PBS. Then first negatively charged peptide was put to 96 well-plate. Congo red having a final concentration of 20 *μ*m (4 *μL*) was added, followed by addition of positively charged peptide (48 *μL* each). Brightfield images were taken immediately with Keyence bz-x710 microscope at defined time points. Finally, ThT assay was performed for aggregation kinetics analysis. Peptides were prepared with the same method as Congo Red assay, but the total peptide volume is adjusted 196 *μL*. Final ThT concentration in peptide solution is 10 *μ*m. Fluorescence measurement were taken immediately after putting positively charged peptide with Ex: 440±10 nm, Em: 480 ±10 nm.

### Peptide Treatment and Viability Experiments

Positively and negatively charged self-assembling peptides were mixed 30 min prior to experiment except where otherwise stated. Viability of the cell lines was measured by using CellTiter-Glo 2.0 Reagent (Promega G9248). Luminescent signal was measured in accordance with the manufacturer’s instructions. Measurements were carried out with BioTek Neo2SM microplate reader and relative viability was calculated. Live-Dead assay was carried out by using Viability/Cytotoxicity Assay kit (BIOTIUM 30002). After peptide treatment, cells were treated with calcein and ethidium homodimer in accordance with the manufacturer’s instructions. Fluorescent images were taken by using Keyence bz-x710 microscope.

### Induction of different cell death modalities

OVCAR-8 cells were treated with 50 *μ*m cisplatin for 24h to induce apoptosis. T75 flask containing OVCAR-8 cells were incubated in a 50°C waterbath for 10 minutes followed by a 24h recovery period for heat induced cytotoxicity. Three cycles of 3 min liquid nitrogen followed by 5 min of 37 °C incubation is used to induce F/T.

### Protein Extraction and Quantification

Lysate proteins: RIPA lysis and extraction buffer (Thermo Fisher Scientific 89900) supplemented with halt protease & phosphatase inhibitor cocktail (Thermo Fisher Scientific 78440) is used to obtain lysate proteins. Supernatant proteins: Acetone precipitation is used for supernatant protein isolation. Briefly, ice-cold acetone was added on supernatants (4:1, v:v), incubated at -20°C for 60 min and centrifuged at 10,000 g for 10 min. Acetone was removed and RIPA buffer is used to resuspend the proteins. Cell lysates were incubated for 15 min on ice and centrifuged at 14,000 g for 15 min at 4°C and supernatants were collected according to the manufacturer’s instructions. BCA protein assay kit (Thermo Fisher Scientific 23225) was used to quantify protein concentrations by measuring the absorbance at 562 nm in accordance with the manufacturer’s instructions.

### Immunoblotting, Reagents

Acrylamide/Bis-acrylamide, 30% solution (Sigma A3699), 1.5 m Tris-HCl, pH 8.8 (Teknova T1588), Tris HCl Buffer 0.5 m solution, sterile pH 6.8 (Bio Basic SD8122), Ammonium persulfate (Sigma A3678), UltraPure 10% SDS (Invitrogen 15553-027), TEMED (Thermo Fisher Scientific 17919), Dithiothreitol (DTT) (BIO-RAD 1610610) Tris Base (Fisher Bioreagents BP152), Glycine (Fisher Bioreagents BP381), 4x Laemmli sample buffer (BIO-RAD 1610747), TWEEN 20 (Sigma P9416), Mini Trans-Blot filter paper (BIO-RAD 1703932), Nitrocellulose Membranes 0.45 *μ*m (BIO-RAD 1620115), Western Blotting Luminol Reagent (Santa Cruz sc-2048).

### Procedure

Protein samples were diluted in Laemmli buffer and boiled for 5 min at 96 ºC. Proteins were then separated by sodium dodecyl sulfate-polyacrylamide gel electrophoresis (SDS-PAGE) gels (8.5% and 15%) and transferred to nitrocellulose membranes. After transfer, the membranes were blocked in 5% milk in Tris Buffered Saline with 0.1% Tween 20 (TBST) or in 5% BSA for phosphorylated antibodies. Blots were then incubated with primary antibodies overnight. Next day blots were washed with TBST and incubated with HRP conjugated secondary antibodies. Lastly, western blotting luminol reagent solution was added on top of the membranes and chemiluminescence signal was detected by Azure c600 Imaging Biosystems.

### LDH Assay

LDH release is measured by Cytoscan-LDH Cytotoxicity Assay (Cat # 786-210). Briefly, media from treated and untreated cells were collected at indicated time points and mixed with reaction mixture and incubated at room temperature. Reaction is stopped by using stop solution and absorbance was measured at 490 nm and 680 nm. Triton-X treatment is used as a 100% LDH release positive control. Lastly, absorbance values at 680nm (background signal) were subtracted from absorbance values at 490nm and relative LDH release was calculated based on the positive control LDH release.

### Immunocytochemistry

Phalloidin staining OVCAR-8 cells were seeded on glass coverslips in a 24 well plate. Cells were treated with [II] for 1-3 and 6h. After each treatment period, cells were washed with 1X PBS and fixed with 4% PFA for 20 min. Then, stained for 30 min with Phalloidin-iFluor 555 (Abcam ab176756) in 1% BSA. After the staining coverslips were mounted in ProLong™ Glass Antifade Mountant with NucBlue™ Stain (Thermo Fisher P36981) and stored in dark until imaging.

### Cytochrome-c staining

OVCAR-8 cells were seeded on glass coverslips in a 24 well plate. Cells were treated with [II] for 1-2-3 and 4h. After each treatment period, cells were washed with 1X PBS and fixed with 4% PFA for 20 min. Then cell membranes were permeabilized by using Digitonin (0.002% in PBS) for 10 min on ice, blocked with 3% BSA in PBS and stained with cytochrome-c antibody in 1% overnight. The next day, samples were washed 3 times with PBS and incubated for 1 h at room temperature with Donkey Anti-Mouse IgG NorthernLights™ NL493-conjugated Antibody (NL009) in 1% BSA. Wells washed with 1% BSA and coverslips were mounted in ProLong™ Glass Antifade Mountant with NucBlue™ Stain (Thermo Fisher P36981) and stored in dark until imaging.

### THP-1 differentiation

THP-1 monocytes were differentiated in the presence of 200 ng/mL phorbol 12-myristate-13-acetate (PMA) for 2 days followed by one day of rest.

### Ethics

Study was carried out accordingly with the recommendations of Guide for the Care and Use of Laboratory Animals from National Institute of Health. Animal procedures were approved by the OU Health Sciences Center (OUHSC) Institutional Animal Care and Use Committee (protocol number 20-059-CHI).

Immunization and Sera Collection Balb/c mice (n=6) were immunized subcutaneously with 5*μg* of HA antigens, 0.5 mM [II] and their combination. Booster was done at D60 and sera were collected at D14, D28 and D75. 4 different influenza antigens; Recombinant HA protein from flu A/Brisbane/59/2007 (H1N1)(NR-28607), HA from influenza A/New Caledonia/20/99 (H1N1) (NR-48873), HA Protein from flu Virus A/St. Petersburg/100/2011 (H1N1) (NR-34588) and Recombinant HA protein from flu A/California/04/2009 (H1N1) (NR-15749) were obtained from BEI Resources. Peptides are mixed for 30 min as explained in the manuscript and mixed with the antigens at room temperature to formulate the vaccines before immunizations.

### ELISA

To analyze the antibody formation, enzyme-linked immunosorbent assay (ELISA) Nunc MaxiSorp^™^ flat-bottom plates (Invitrogen 44-2404-21) were coated with 2.5 *μg*/mL with the mixture of four antigens in phosphate coating buffer (0.1 M Na2HPO4 in deionized water, pH 9.0) overnight at 4^*°*^C. Next day, plates were blocked with 1% bovine serum albumin in phosphate buffered saline-Tween (1X PBS, 0.05% Tween) for 2h at room temperature. Plates were then washed 4x with PBS-T and incubated overnight at 4^*°*^C with serially diluted sera collected from mice in PBS-T. The next day, wells were washed 4x with PBS-T and incubated for 1h at room tepmerature either with HRP-IgG1 (SouthernBiotech 1070-05) (1:4,000) or HRP-IgG2a (SouthernBiotech 1080-05) (1:4000). Wells were subsequently washed 4x with PBS-T and developed with 2,2-azinobis(3ethylbenzthiazolinesulfonic acid) (ABTS) (VWR 95059-146) substrate for 5 min at room temperature. At the end of the incubation, reaction was stopped with 10% SDS solution. Absorbance was measured at 405 nm to determine endpoint antibody titers.

## Notes

### Competing Interest Statement

The authors have declared no competing interest.

